# Whole Body PBPK Model of Nasal Naloxone Administration to Measure Repeat Dosing Requirements During Fentanyl Overdose

**DOI:** 10.1101/2023.04.24.538032

**Authors:** Austin Baird, Steven A. White, Rishi Das, Nathan Tatum, Erika K Bisgaard

## Abstract

Opioid use in the United States and abroad is an endemic part of culture with yearly increases in overdose rates and deaths. As rates of overdose incidence increases, the use of the safe and effective reversal agent, naloxone, in the form of a nasal rescue spray is being fielded and used by emergency medical technicians (EMTs) at a greater and greater rate. Despite advances in deployment of these rescue products, deaths are continuing to increase. There is evidence that repeated dosing of a naloxone nasal spray (such as Narcan) is becoming more common due to the amount and type of opiate being abused. Despite the benefits of naloxone related to opioid reversals, we lack repeated dosing guidelines as a function of opiate and amount the patient has taken. Goal directed dosing is promising, where respiratory markers are being used as an indication of the patient recovery but require time and understanding by the EMT. We construct a whole-body model of the pharmacokinetics and dynamics of an opiate, fentanyl on respiratory depression. We then construct a model of nasal deposition and administration of naloxone to investigate repeat dosing requirements for large overdoses. We demonstrate that naloxone is highly effective at reversing respiratory symptoms of the patient and recommend dosing requirements as a function of opiate amount administered. By designing the model to include circulation and respiration we investigate physiological markers that may be used in goal directed therapy rescue treatments.

## 1 Introduction

The opioid epidemic continues to kill countless people across the globe. In the United states alone 47,600 people have died in 2018 (Wilson et al., 2020), this reflects an increase of 5,100 since 2015 (Scholl et al., 2018). Due to the amount of people dying due to opioid related overdose there has been a concerted and large effort to equip all emergency medical technicians to carry nasal naloxone as an effective reversal agent (Sumner et al., 2016). This equipment and subsequent increase in naloxone administration has been directly shown to decrease opioid related deaths in controlled settings and specific regions (Kerr et al., 2009, 2008; Rando et al., 2015). Despite the efforts to properly equip first responders with the appropriate reversal agents, opioid-related deaths are still shockingly high and are showing a marked increase in countries around the world (Amsterdam et al., 2021). Throughout this progression of deaths there has been an associated increase of naloxone administration, with a 75% increase seen in between the years 2021-2016 (Cash et al., 2018). This increase in administration mirrors the increase in opioid associated overdose deaths during that period. The coupling of naloxone administration to opioid associated overdose deaths is hypothesized to be related to more synthetic opiate use (Zibbell et al., 2019), more opiate use in the general population (Seth et al., 2018), inadequacies in naloxone reversal dosing (Moss and Carlo, 2019), or compounding physiologic factors that due to the type of opioid being used (Torralva and Janowsky, 2019).

Synthetic opiates have become prominent in America over the past decade, particularly illicitly manufactured fentanyl (Zibbell et al., 2019). During this time increase in synthetic opiate deaths have outpaced traditional prescription opioids (Seth et al., 2018). The nature of the pandemic has evolved as the type and potency of the drug has changed. This has placed a burden on first responders to administer more naloxone without an understanding of the differences between overdose related to fentanyl and morphine with consensus that naloxone reversal dosing is inadequate to properly rescue many patients (Moss and Carlo, 2019). Studies have shown that overdose due to fentanyl may require more naloxone than what is currently in a single dose of Narcan (the most common form of reversal agent equipped to EMTs around the country) (Faul et al., 2017). This is despite many counties still administering the recommended amount of naloxone despite the evolving nature of opioid epidemic (Bell et al., 2019). Indeed, in response to this suffering much of the United States opioid education and harm reduction strategy includes distributing naloxone directly to citizens. Although promising, there is evidence that this response isn’t sufficient due to the higher naloxone dosing requirements for most illicitly manufactured fentanyl derivatives (Kim et al., 2019). Due to the evolving nature of the issue, more understanding regarding dosing requirements for different types of opiates is needed. This is particularly true to the for common EMT nasal spray form of naloxone, Narcan 2 and 4mg doses.

To quantify and analyze the dose requirements and needs of first responders in relation to nasal naloxone administration, we develop a whole-body physiological model that can properly capture different dosing amounts, physiological respiratory depression, and model the interaction between an opiate and naloxone in the body. To date various models have constructed nasal deposition and absorption models, with much of the work focused on 3D models of deposition (Foo et al., 2018; Frank et al., 2012). Other work has been done to understand pharmacokinetic profiles of a drug in the blood after nasal administration (Corley and McMartin, 2005; Dave et al., 2022; Gonda and Gipps, 1990; Rygg et al., 2016). Generally, these models consider breaking the nasal cavity down into a series of compartments with transfer rates corresponding to the drug being considered or the transport mechanism being modeled. Despite this, there are no models that consider dynamic interactions between the patient physiology, nervous system response, and repeated naloxone dose and interaction with different opiates.

The software architecture of BioGears allows for refined questions to be asked regarding naloxone administered as a nasal spray and its relation to the physiology of the patient experiencing opioid overdose symptoms. Here we will describe a general pharmacokinetic model of opiates, using the BioGears engine, and a kinetic model of naloxone administered via a nasal spray device. We will also describe a model of competitive mu-receptor drug interaction between the opiate and naloxone. We consider the opioid respiratory depression via a pharmacodynamic model that is coupled to a whole-body nervous system that captures large scale regulatory responses in the body. Using these models, we will investigate dosing requirement differences apparent between fentanyl and morphine and investigate how body type plays a role in this calculation. We also present a model of the pharmacodynamics of these drugs and their physiological impact on the patient. We consider the respiratory depression associated with opioid overdose and hope to answer some of the following questions, pertinent for first responders: when to administer naloxone, how long to observe the patient before leaving, and when to administer a possible second and subsequent doses?

## 2 Materials and Methods

The BioGears Engine fluid transport model constructs the analogous circuit model to characterize the fluid dynamics of the cardiopulmonary system. We characterize the fluid and gas flows, pressures, and volumes by their electrical equivalents; current, voltage and capacitance respectively. Circuit analogues have been used frequently in physiology modeling, usually to model cardiac pumping dynamics using a windkessel model (Beard et al., 2005; Westerhof et al., 2009, 1969). BioGears expands this construction by using a series of connected three element windkessels and other more sophisticated circuit architectures, to model blood flow in each organ and in major vessels (Baird et al., 2020a). The software architecture of BioGears constructs objects representing possible circuit elements, allowing for organs, like the brain and kidney, to have larger, more sophisticated circuit structures, to increase fidelity, Figure 1.

**Figure 1.**
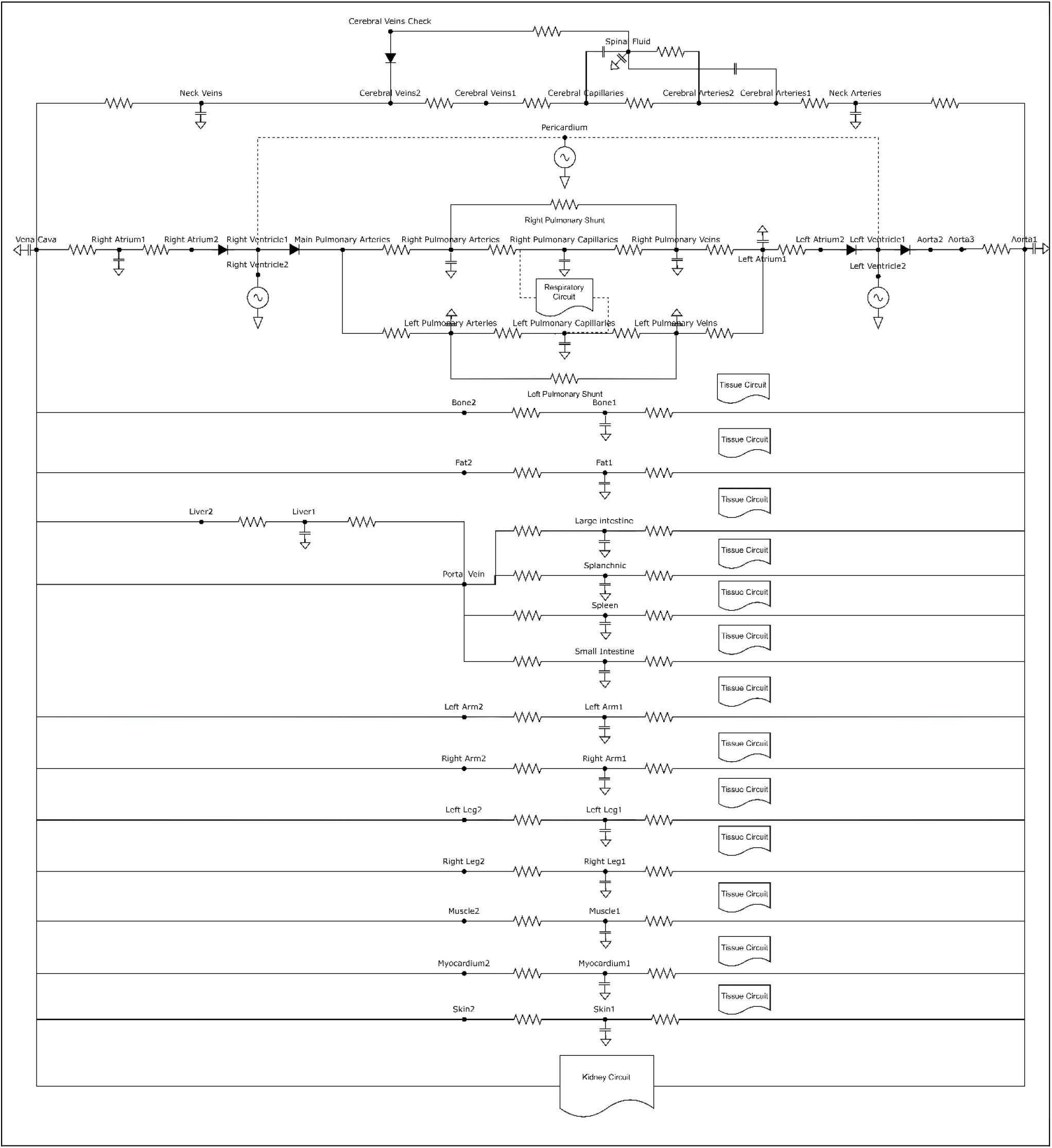
Circuit diagram of the BioGears circulatory system. The additional boxes denote broken out circuit structure for each of the noted biological system, the renal, tissue, and respiratory (gas) systems have their own separate circuits that connect to the vasculature through various junctions not shown here.

Pumping of the heart occurs by adjusting the compliance of the cardiac windkessel, effectively simulating contraction of the heart chambers. The BioGears respiratory system contains models defining gas exchange and diffusion. Collectively, these circuits constitute a lumped parameter, or “zero-dimensional,” system. That is, because no spatial component exists, the pressure, volumes, and flows calculated on each circuit represent values averaged (lumped) by each organ, vessel, and tissue component. Such an approach is appropriate for a model of this size considering the computational cost incurred by increasing fidelity (Shi et al., 2011). Higher dimensional models require numerical solutions of either the full Navier-Stokes equations, or a simplification assuming, for instance, radial symmetry or low Reynolds number (Batchelor, 1970; Peskin, 2002). We choose this vasculature and cardiopulmonary fluid transport model to simulate and investigate naloxone reversal over hours post opioid overdose. Higher fidelity models would not be sufficient for the time scales considered for this research.

The BioGears physiology Engine organizes the data generated by the circuit structure and the various other connected models, representing the state of the patient, hierarchically into compartments. Top-level compartments generally represent organs or systems, with sub-compartments representing the vascular, tissue, extracellular, and intracellular spaces. The compartment structure is computationally organized as a graph to increase data processing speed. This graph is then leveraged to simulate the movement of substances in the body. Once the circuit is calculated and flow rates computed, the compartment graph then moves substances as a function of these flow rates and checks for mass conservation. Some transport connections require a more refined model, such as diffusion from the alveolar gas compartment into the alveolar capillaries. These connections are computed separately after the general substance movement is resolved.

The engine implements a Compartment Manager class that organizes and keeps track of compartment graphs and hierarchal mappings. Each simulation cycle, the state of the circuit is solved using a modified nodal analysis algorithm (Ho et al., 1975). BioGears leverages the third party numerical software Eigen 3 to compute the matrix inversion using a sparse LU factorization (Guennebaud et al., 2010; Saad, 1994). Substance flux, diffusion, and mass transfer is computed from the flow rates and tissue properties using the compartment graph and manager class. Various system and subsystem models are computed using a basic forward Euler scheme, for non-stiff equations.

Custom implementation of an improved Euler method using two functional evaluations per time step is used to compute differential equations with more rapid dynamics. Each system implements its own class, providing the user the ability to directly implement methods specific to a certain organ. The system classes implemented in BioGears contain all the code used described for this paper, specifically the drugs, respiratory, and nervous class files. We omit other model descriptions here for brevity but refer the reader to the open-source documentation for a full description.

The circuit and graph constructs are the backbone of the physiology engine with numerous physiological models (such as a robust nervous system) developed on top of, and often influencing, this backbone (McDaniel et al., 2019b; McDaniel and Baird, 2019). These models include: a nervous system with baroreceptor and chemoreceptor feedback that modifies cardiovascular and respiratory activity; an active transport model that maintains ionic gradients across intracellular and extracellular compartments; a physiologically-based pharmacokinetic/pharmacodynamic (PBPK/PD) model that tracks the concentration-effect profile of numerous drugs (McDaniel et al., 2019a); a gastrointestinal model that determines rates of nutrient digestion and oral drug absorption; a renal feedback model that regulates urine production and substance filtration and reabsorption; and a metabolic consumption and production method that determines the energy demands of each organ. In addition, numerous injury models have been developed to influence the state of the patient ranging from acute hemorrhage to diabetes. Taken all together this platform provides the unique opportunity to not only investigate the kinetics of the drug in the body but also how opioid overdose impacts the physiology and oxygen transport in the body. We will leverage this existing cardiopulmonary and nervous implementation to investigate naloxone reversal timing and dosing requirements for different opioid overdose situations.

## 2.1 Nasal Administration Model

### 2.1 1 Pharmacokinetic and Transport Model

Circulation of substances in BioGears is driven by a lumped model of the cardiopulmonary system, which is based on prior physiological modeling. Drug administration models the substance mass in the appropriate compartment, depending on the administration route: intravenous, intraarterial, nasal, oral, intramuscular, and inhaled dose. Oral, nasal, and inhaled dose each consider deposition, digestion, and absorption into the appropriate tissue. For example, oral doses are absorbed into the vascular system via the small intestine compartment and inhaled doses are deposited into the lung tissue and absorbed into circulation via the alveolar compartment. For nasal naloxone we consider the compartmental structure of the nasal cavity with appropriate absorption metrics (section 3.1). Upon entering the circulation, we use perfusion limited diffusion to deposit the drug in the various tissue compartments as it circulates. We expand upon prior work to compute the appropriate partition coefficient for a given tissue and substance (Rodgers et al., 2005; Rodgers and Rowland, 2006).

Hence, for a given drug in circulation we compute the change in mass over time as the concentration gradient for a given blood and tissue compartment, scaled by a computed partition coefficient:

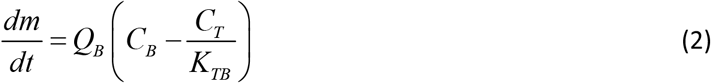

Given a substance’s physiochemical properties, table 1, we can compute the partition coefficient, as:

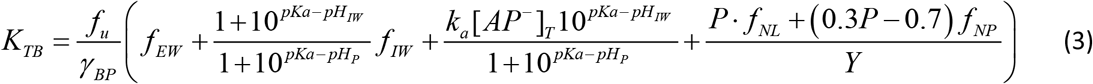

For strong bases. Here we note that *f*_*u*_ scales the classical formulation from plasma to blood. Similarly, for acids, neutral compounds, and weak bases, we have

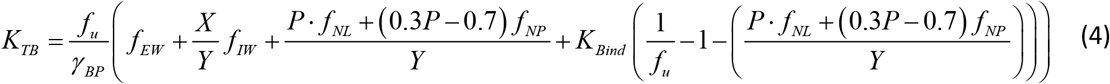

And for zwitter ions we have

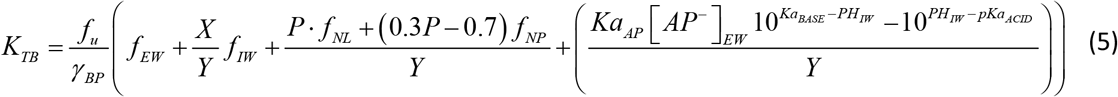

Here the term [*AP*^−^]_*EW*_ is the concentration of acidic phospholipids in the extracellular water and *Ka*_*AP*_ is the association constant for a drug for acidic phospholipids. These three terms allow for computation of a given partition coefficient for a specific tissue and drug combination. The parameters X and Y depend on the state of the drug as follows:

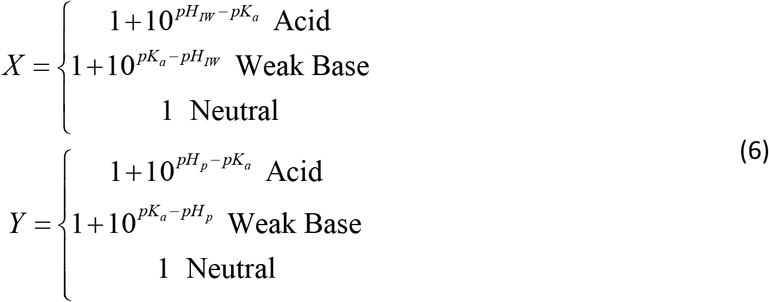

Parameters pH_p_ and pH_IW_ represent the pH of the plasma and intracellular water, respectively. Each parameter is computed at each time step in the simulation and will respond to changes in pH during a simulation if certain injuries present themselves to the patient. For this application they are effectively constant.

**Table 1.**
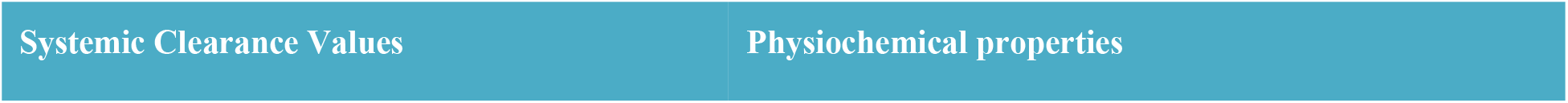
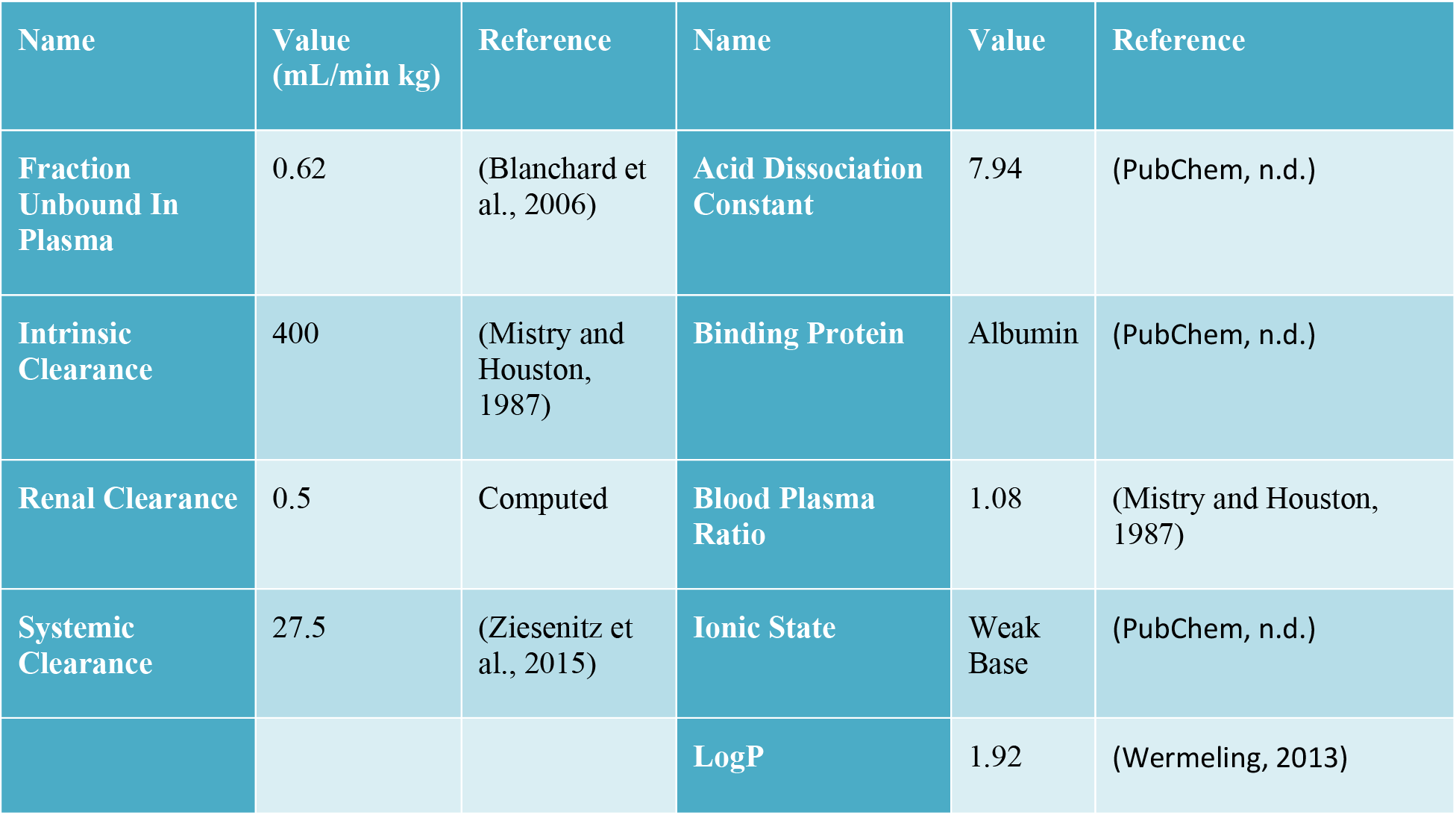
Clearance and physiochemical properties of naloxone. Some parameters have small deviations from reported literature but are generally within the error deviation reported. Changes are due to matching concentration profiles of naloxone in the blood.

| Name | Value<br>(mL/min kg) | Reference | Name | Value | Reference |
| --- | --- | --- | --- | --- | --- |
| Fraction Unbound In Plasma | 0.62 | (Blanchard et al., 2006) | Acid Dissociation Constant | 7.94 | (PubChem, n.d.) |
| Intrinsic Clearance | 400 | (Mistry and Houston, 1987) | Binding Protein | Albumin | (PubChem, n.d.) |
| Renal Clearance | 0.5 | Computed | Blood Plasma Ratio | 1.08 | (Mistry and Houston, 1987) |
| Systemic Clearance | 27.5 | (Ziesenitz et al., 2015) | Ionic State | Weak Base | (PubChem, n.d.) |
|  |  |  | LogP | 1.92 | (Wermeling, 2013) |

Clearance of the drug from the blood stream occurs by hepatic and renal elimination. Here we define hepatic clearance by

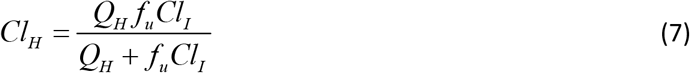

Here Q_H_ represents the volumetric rate of blood supplied to the liver by the portal vein. In BioGears the small intestine, large intestine and splanchnic compartments send blood flow into the portal vein. Cl_I_ represents intrinsic clearance and is normalized by the patient body weight. We then compute the mass of drug metabolized and removed from the liver compartment at each time step:

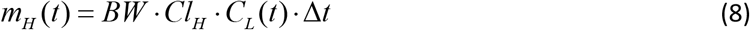

We define Cl_L_(t) as the concentration of the drug in the liver vasculature at time t. Other clearance elimination mechanisms are supported by Biogears including renal, fecal, and systemic and are computed similarly.

We compute the pharmacodymics of an opiate circulating in the body by computing an effect site concentration of the drug:

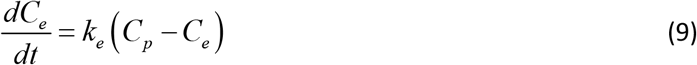

This effectively models the transport of the drug from the systemic circulation to the site of action, or binding. Here ke is a first order rate constant buffering the binding of the drug to the effect site, Cp and Ce are the concentration of the drug in the plasma and the effect site. We model effect of the drug by modeling the interaction between an opiate, such as morphine or fentanyl, with its antagonist, naloxone, with the following functional relationship:

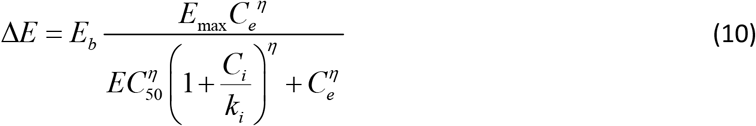

Here Eb is the baseline of the given physiological effect the drug is exerting on the patient, such as cardiovascular or pulmonary effects. Emax is the maximal possible effect, EC50 is the half maximal effect, and lastly Ci and ki are the inhibitor concentration and constant. For example, in the case of morphine, the inhibition concentration is the concentration of naloxone in the effect compartment. There are numerous possible effects that can be mapped to a specific drug in the BioGears physiology engine, but we will omit those effects for brevity. A full list can be found in the documentation (n.d.). The way the drug is administered will affect the concentration profile in the blood and our model will consider diffusion and release of the drug through the various nasal compartments to enter the bloodstream.

### 2.1.2 Nasal deposition model

Nasal drug administration offers an effective alternative drug delivery route by utilizing the large surface area of the nasal canals and the porous endothelial membrane which lines them. The unique physiology of the nasal canal offers a vasculature space which clears drugs rapidly (Türker et al., 2004). Much like the intravenous or intramuscular delivery methods, nasal drug delivery skips the first pass effect of the metabolism (Graff and Pollack, 2005). However, it provides similar benefits to oral dosing in that the nasal canal is a noninvasive pathway for rapid systemic drug absorption. While the narrow and nonlinear structure of the nasal canal can cause some challenges in drug retention, the nasal delivery method provides optimal opportunity for drug exposure in a readily accessible system. In addition, most emergency rescue equipment is home to at least one form of nasal rescue spray, we will focus this paper on nasal delivery methods. We note, that the BioGears physiology engine also supports intramuscular, intravenous, and intraartial administrations in addition to the nasal route described here.

We define our nasal drug delivery model by building upon existing pharmacokinetic and transport models (Dave et al., 2022; Foo et al., 2018; Gonda and Gipps, 1990). Following drug release, we track penetration and permeation through the nasal mucosa (modeled by anterior nose and posterior nose compartments). By tracking the amount of drug released through each layer. We construct a series of first order differential equations to solve for the drug quantities excreted through the gastrointestinal tract and the amount absorbed into systemic circulation. In BioGears, drug deposited into systemic circulation is then be managed by the substance transport and existing pharmacokinetics and dynamics models, 2.1.1.

We relate the transition of a drug administered through the nasal cavity to its transport through the anterior (A), posterior (P) and gastrointestinal (G) compartments:

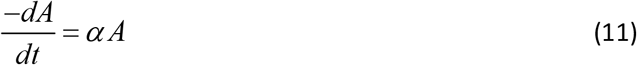

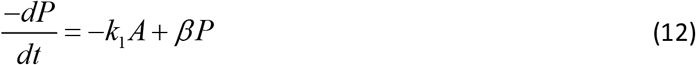

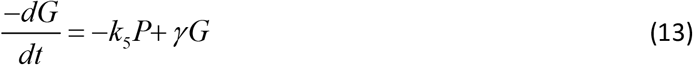

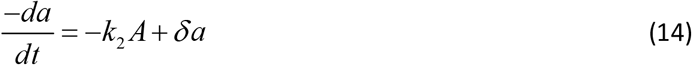

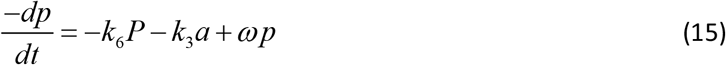

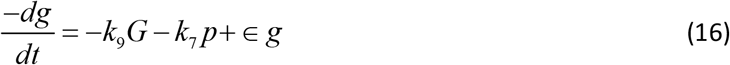

Where:

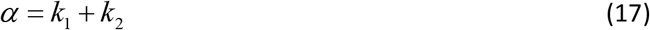

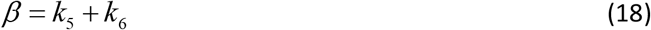

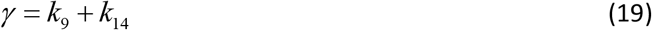

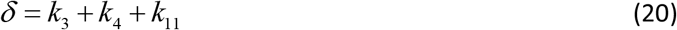

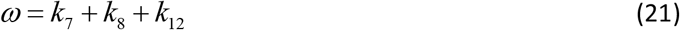

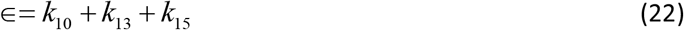

To produce the rate of drug input into systemic circulation:

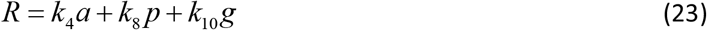

We note that the difference between upper case and lower casing for this model delineates between drug unreleased in a carrier solution and drug released from that carrier. In BioGears, we update our vector of released and unreleased nasal drug quantities in each compartment (A, P, G, a, p, g). We solve this system of equations and multiply by the timestep to get the amount of drug mass to be moved into our vasculature compartments. We note that we only consider the mass of the drug at each compartment in the nasal canal. We do not consider the changes in deposition due to concentration of the administered spray, as there is some evidence that concentration of the spray does have an effect on the kinetics of the administered dose (McDonald et al., 2018).

Table 2. First order rate constant (1/s) used in the ODE’s for modeling nasal drug administration. Rate constants taken from (Gonda and Gipps, 1990; Graff et al., 2005). Minor changes have been made in order to fit the model to the specific naloxone pharmacokinetic profiles of experimental data, whereas this model was first produced as a generic model of nasal administration.

**Table 2.** Symbols and their associated meanings for the nasal compartment ODE model.

| Symbol | Description | Symbol | Description |
| --- | --- | --- | --- |
| **A** | Amount of drug in the anterior (nonciliated) nose (mg) | **p** | Amount of drug released in the posterior (ciliated) nose (mg) |
| **P** | Amount of drug in the posterior (ciliated) nose (mg) | **g** | Amount of drug released in the gastrointestinal tract (mg) |
| **G** | Amount of drug in the gastrointestinal tract (mg) | **R** | Total rate of drug input into systemic circulation (mg/s) |
| **a** | Amount of drug released in the anterior (nonciliated) nose (mg) |  |  |

**Table 3.** Kinetic rate parameters used for the nasal administration model.

| Symbol | Value | Symbol | Value |
| --- | --- | --- | --- |
| $k_1$ | 0.00001736 | $k_9$ | 1000000 |
| $k_2$ | 1000000 | $k_{10}$ | 0.000000027 |
| $k_3$ | 0.00001736 | $k_{11}$ | 0.00001 |
| $k_4$ | 0.000173 | $k_{12}$ | 0.00001 |
| $k_5$ | 0.0011575 | $k_{13}$ | 0.00001 |
| $k_6$ | 1000000 | $k_{14}$ | 0.0000278 |
| $k_7$ | 0.0011575 | $k_{15}$ | 0.0000278 |
| $k_8$ | 0.0000260 | | |

### 2.2 Central and Peripheral Nervous System Model

Perturbations to oxygenation in the BioGears system are regulated and responded to through a robust nervous system model. Efferent and Afferent responses are managed and weighted in the central nervous system model. The afferent baroreceptor, chemoreceptor and pulmonary stretch receptors are computed and collected by the central nervous system which weights these responses and generates the efferent response on the appropriate system in BioGears. The chemoreceptor control model in BioGears is based upon prior model development that considered hypoxic conditions on the nervous system (Cheng et al., 2016; Magosso and Ursino, 2001; Ursino and Magosso, 2004, 2000), we connect these models to our model of the blood gas binding, respiratory driver function, pharmacodynamic, and transport models in order to elicit a patient response depending on a variety of perturbations to the patient state. The chemoreceptors are modeled as two signals: a central and peripheral response and their contributions are independent and additive, for the frequency adjustments to ventilation we have:

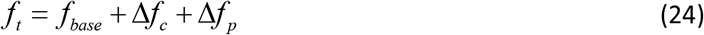

Similarly for the driver pressure response:

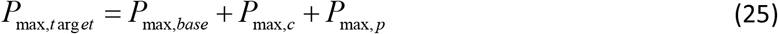

We define each of these as respiratory effects *E*_*c, p*_ where c denotes central effects and p denotes the peripheral chemoreceptor response. Using this definition, we denote the evolution of this response to be a function of arterial carbon dioxide partial pressure 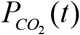 (*t*) changes against a normal homeostatic set point 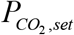:

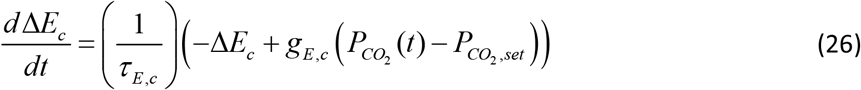

Here the effects *E* = { *f, P*} are respiratory driver frequency *f* and amplitude *P* with associated time constant τ_*E, c*_ and controller gain *g*_*E, c*_. We then define the peripheral feedback in a similar way to equation 25, but here the response is a function of the combines CO2 and O2 perturbations. As previously the effector equation computes a response for driver frequency and amplitude and is defined as:

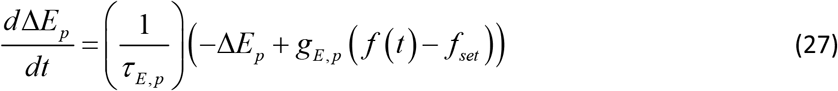

Here, *f* (*t*) denotes the firing rate of the response and is a nonlinear sigmoid function of CO2 and O2:

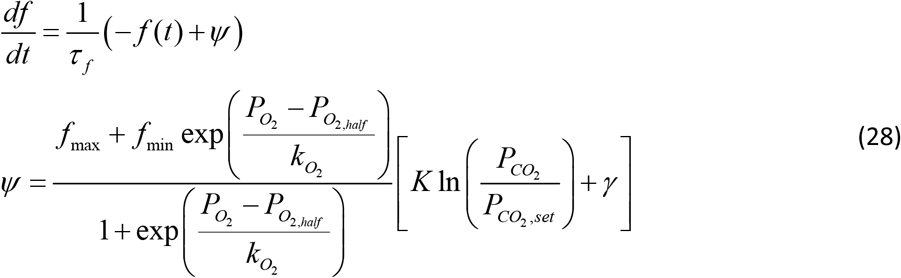

Here 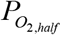 is the half max and 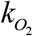 is the slope of the sigmoid response. K and γ are tuning parameters used to maintain a steady state response under normal conditions. Values for the parameters are mostly taken from prior studies, (Magosso and Ursino, 2001), or are tuned to meet validation data under various patient hypoxic states due to different injuries. The central nervous system then takes the chemoreceptor effects and weights them to compute an efferent response.

Opioids act against this central nervous by binding to the mu receptor, buffering the system’s ability to control hypoxic conditions.

## 3. Results

### 3.1 Pharmacokinetic Profiles

To test the proper dosing of naloxone needed to reverse extreme overdoses involving a synthetic opioid, such as fentanyl, we validate the nasal model of naloxone. We collected data for 2, 4, 8, and 16mg doses of naloxone and their associated venous plasma concentrations. The low dose concentration data was collect from (McDonald et al., 2018). Their study randomized 38, healthy male and female patients.

With 19 blood samples taken over a 2-hour study period. We do note that the study population trended male, consisting of 71 % of the study participants. Standard deviations were reported by the authors and that data extracted for this comparative study. The experimental study also collected intramuscular, intravenous and 1 mg intranasal dose, but the results were omitted for this modeling study.

By hypothesizing that the naloxone required to resuscitate a patient who has been given an extreme dose of fentanyl we also wanted to investigate the pharmacokinetics of naloxone doses above the average dose administered by a single Narcan nasal spray. To this end, we collected 8mg and 16mg naloxone nasal dose data from (Mundin et al., 2017) to test the model against. No additional parameter fits were performed to evaluate these higher doses. The study enrolled 12 subjects, 5 male and 7 female, for the study with varying age, weight, and height. The naloxone nasal dose was formulated in the lab from a powdered dose mixed with sodium citrate and sodium chloride and atomized in a metered spray device. Blood was sampled at 10 distinct points throughout the six-hour study period with standard deviations computed for the population.

The nasal naloxone model presented here was fit directly to the 2mg nasal dose (training) with the other doses presented here (4, 8, and 16 mg) used as validation (test). No kinetic parameters were changed after the fit to the 2 mg dose. As the naloxone circulated in the blood (represented here as a lumped parameter circuit system). Deposition and clearance were performed through computation of the partition coefficient for a given tissue compartment, Equation 2. Physiochemical properties of naloxone were used to compute the partition coefficient and none of these parameters were altered during the dose study, Table 1.

To analyze the validity of using this model to study reversal and resuscitation efforts in the overdose patient, we first compared the validated data to the simulated data through the root mean square error (RSME). We note that root mean square error scales with the size of the naloxone dose (non-normalized), so to aid in understanding of the errors in the model we also compute an average percentage that corresponds to the percentage that the simulated data is different from the validation data (on average), Table 4. We also note that because the 8mg and 16mg data was collected from a different source, there were more data points available and a slightly different experimental protocol for the participants. This shouldn’t impact the RSME as additional data points just contribute to the average distance incrementally, not in a significant way.

**Table 4.** Root mean square error of simulated naloxone kinetics in the body after nasal administration. Validation data was used to fit kinetic model for the 2mg dose. The error term is an average vertical distance between the validation data and simulated data. The percentage is computed by scaling the error term by the maximum naloxone concentration value.

| Naloxone Dose (mg) | Root Mean Square Error | As a percentage of peak concentration |
| --- | --- | --- |
| 2 | 0.324 | 12 |
| 4 | 0.806 | 14 |
| 8 | 1.08 | 11 |
| 16 | 2.36 | 17 |

We note that the simulated data matches rate of onset and long-term dynamics quite well, with much of the error being generated in the peak concentration profiles, Figure 2 and Figure 3. The 16 mg dose displays the largest deviation from experimental data with a significant overshooting of the peak reported naloxone concentrations. We do note that the 16 mg experiment performed does display the largest reported standard deviations, especially early in the first few data points. In general, peak profiles were underestimates in the model although all simulated data stayed under 20% deviation throughout each experiment.

**Figure 2.**
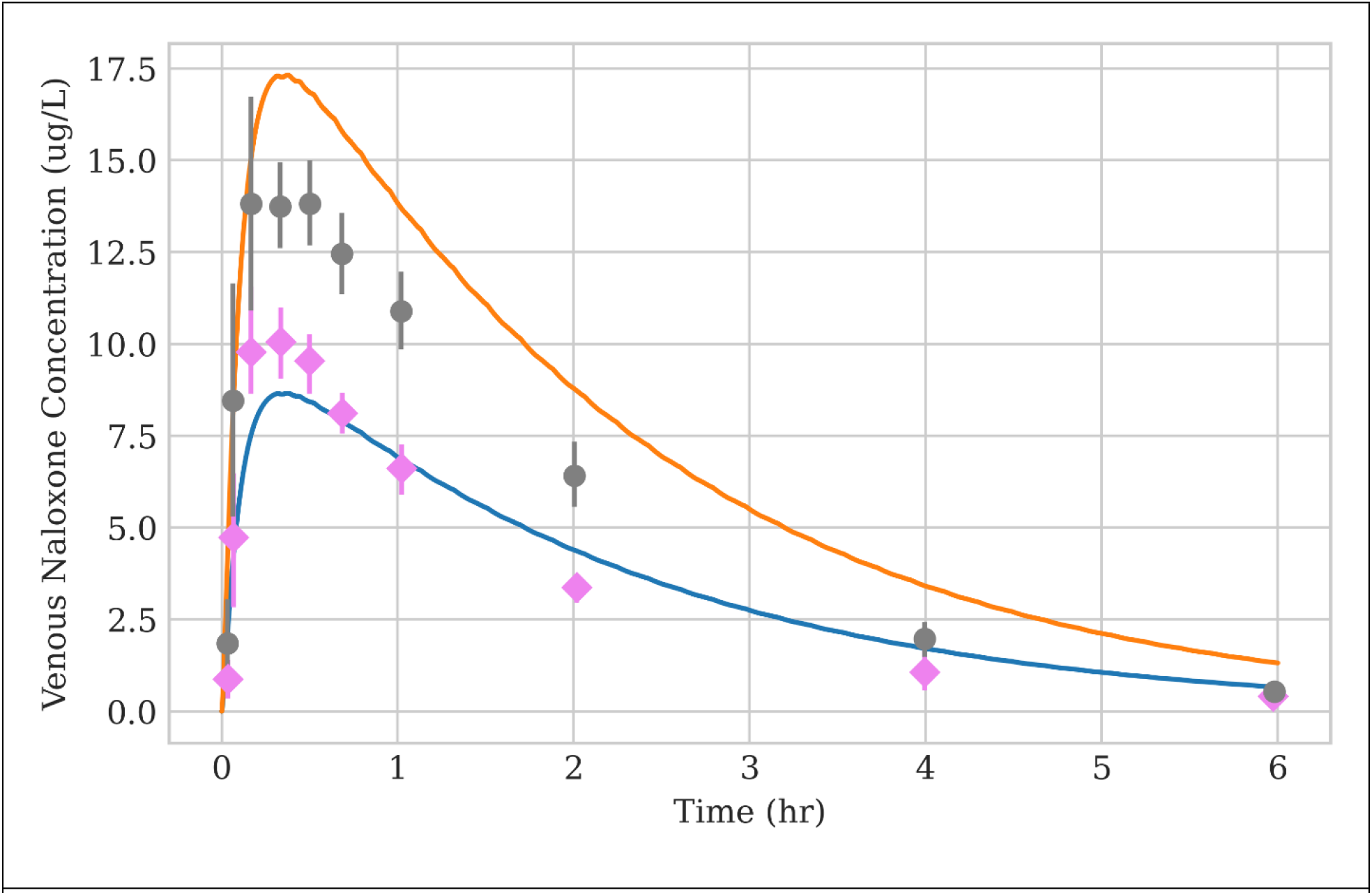
6-hour simulation of naloxone for 16 mg (orange) and 8 mg (blue) metered nasal spray doses. 16 mg simulated data clearly overshoots the peak profile of the experimental data. Both simulations match the early onset growth and long-term dynamics well.

**Figure 3.**
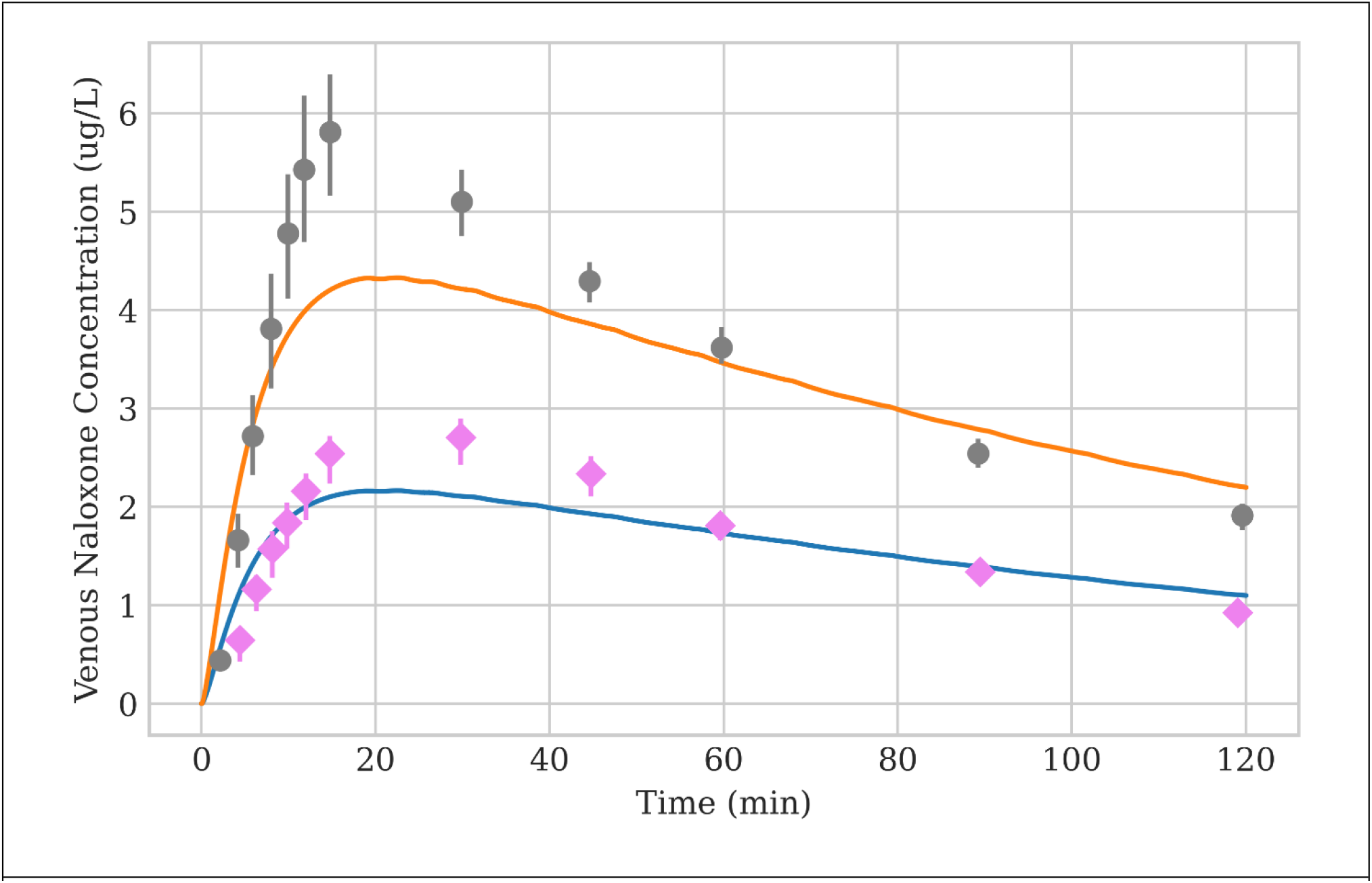
2-hour simulation of nasal naloxone dose for 4 (orange) and 2 (blue) mg doses. The peak venous concentration and long trend dynamics follow experimental data well, with minor peak concentration differences for each dose.

### 3.2 Naloxone Treatment Analysis

To begin testing the naloxone administration amount needed for fentanyl overdose reversal we first created an algorithmic scenario centered around the patient’s oxygen saturation (SbO2). Assuming first responders would be able to access this piece of data, we built a repeat administration scenario to properly recover the SbO2 over the length of the scenario. To begin the scenario, fentanyl was administered, followed by a two-minute waiting period. To assess the reversal dosing requirements, we incremented this initial fentanyl dose from 0.2 mg up to 1.9 mg, generally considered a very high dose for recreational use. We note that if the patient did not receive any nasal naloxone dosing, the patient would die for doses of fentanyl over 1.5mg.

To simulate the transit time of the emergency care provider we waited 2 minutes post opioid dose before administering naloxone. Once that two-minute wait period has expired we administered an initial 2mg rescue dose naloxone. After the initial rescue does was administered, we continuously monitored the patient’s SbO2 and at each 50 second increment, evaluated whether it was above a certain threshold value. We chose a value of 85% as a level of concern clinically. We note that generally a value below 90% would be considered hypoxic but did not want to overcompensate our dosing amounts by administering naloxone in the latter stages of the patient recovery. More complex goal directed therapies are possibly within the physiology engine, but we wanted to only understand the naloxone administration requirements as the sole rescue intervention. Other resuscitative interventions could be considered in the future, such as ventilation, bag valve mask, or supplemental oxygen. Additionally, the resuscitation scenario could have longer observation periods in between repeat does which may alter the data presented in this manuscript. We note that the end-of-life thresholds for the physiology engine are rather crude and would need to be updated to firmly observe death in a patient from asphyxiation. All data was generated from a single C++ program using the BioGears engine, (Baird et al., 2020b). The HowTo example file is open source in the core repository on GitHub.

### 3.2.1 Respiratory Function

Respiratory depression due to fentanyl administration is produced through binding of the mu-opioid receptors at specific sites in the central nervous system. We simulate this action by reducing the nervous system function in the physiology engine as a function of fentanyl blood concentration. This function in turn, drives the respiratory driver in the physiology engine. Central nervous depression direction depresses respiratory cadence of the patient, thus inducing a hypoxic state in the patient, Figure 5. Respiratory rate is properly depressed as a function of the initial dose of the patient, with mu-opioid receptor saturation incurred at doses above 1.5 mg. Number of breaths per minute in the patient drop to 2 from a baseline rate of approximately 16. Oxygen saturation in the physiology engine is computed as a function of blood-gas binding and oxygen diffusion through the alveolar compartment.

Because respiration rate is so dramatically reduced oxygen saturation rates drop over the scenario duration. Recovery for large doses over 1mg takes much longer than for lower doses of initial opioid administration.

Oscillations seen in the oxygen saturation curves are due to respiratory function and circulatory oscillations seen in the models, as well as delayed interaction with the nervous system model. We note that a common patient response to opioids is irregular breath rhythm and tidal volumes (Bouillon et al., 2003; Lumsden, 1923; Pattinson, 2008). The model presented here does show some minor variability in the respiration rate, although not consistent with the pronounced variability seen in many patients. The tidal volume, Figure 9, does display much more erratic variability, possibly due to the pronounced role of the peripheral nervous system response and associated changes in driver pressure during the simulation. Future work could potentially focus on modeling the coupling of the two primary oscillators present that regulate respiration frequency: pre-Botzinger complex and the nearby retro-trapezoid and parafacial respiratory group, (Del Negro et al., 2018). There is evidence to show that opioid act on only one of these, causing disruptions in the rhythm generated by the dynamic coupling of these oscillators. Finer representation of these neurological actuators and the influence of opioid receptor binding would be an excellent iterative advancement of the models presented here.

### 3.2.2 Cardiovascular Function

The cardiovascular model is regulated by the baroreceptors, which work to regulate elastance during a cardiac cycle. Although not affected by the opioid directly the cardiac output is seen to increase for large opiate use scenarios. This corresponds to the decrease in blood pressure (seen by the decrease in overall systemic resistance) and the slight reduction in heart rate during the early stages of the fentanyl progression. Cardiac output in the physiology engine is computed to be the stroke volume multiplied by heart rate. Even though the heart rate decreases slightly for high levels of opioid use, the stroke volume increases due to lower blood pressure. Stroke volume and most systemic cardiovascular effects are unchanged for opiate doses below the 1 mg threshold. In addition, as oxygen is depleted due to the respiratory depression, the nervous system adjusts the resistance into the myocardium increase fluid flow into the heart muscle tissue to maintain oxygenation.

Fentanyl pharmacodynamics provide some depression to the main drivers of the circulation, namely heart rate and blood pressure. Although small, these changes can be seen for opioid doses over 1 mg, Figure 7. As the opiate is cleared in the body and the respiratory depression is reversed with naloxone, the nervous system can compensate for the lack of oxygen in the blood by spiking the heart rate. We note that this increase is moderate and is an artifact of the injury that the patient has just sustained during the overdose episode. Relaxation of the heart and the vascular resistance is seen as the patient fully recovers at the end of the episode.

### 3.2.3 Nervous System Function

The nervous system in the physiology engine is modeled to provide feedback to the body during times of perturbations from normal homeostatic physiological values. These types of perturbations can occur globally, oxygen partial pressure changes and blood pressure changes, or locally, drops in renal artery perfusion. Local changes are handled through autoregulatory relationships that work to preserve the perfusion to a given region, such as the brain or kidney. During an opioid overdose the patient oxygen partial pressure is perturbed in a significant way, leading to hypoxia. The acting mechanism by which is occurs is through fentanyl depressing the central nervous system. This depression leads to a decreased frequency of respiratory function. The peripheral nervous system is allowed to respond accordingly and the difference between the two responses can be seen in Figure 8. The central frequency can properly respond to the respiratory depression and associated reduction in oxygen partial pressure in the blood after the naloxone can properly reverse the opioid central nervous system binding.

Tidal volume properly responds to the reduction in oxygen in the blood by increasing the amount of volume per breath. This is achieved through increases in the peripheral pressure changes which alter the respiratory diver pressure. This pressure difference creates a larger negative potential for the lungs to fill during a single cycle, Figure 9. We note that the tidal volume generally should decrease for this amount of fentanyl administration with other research suggesting decreased volumes during increased opioid exposure, (Pattinson, 2008). Other opioids have been shown to not decrease tidal volumes in various studies, (Hill et al., 2020) leading to unique respiratory responses depending on the type of opiate the patient has been exposed to. The qualitative issues shown here can be mitigated in future work by including a term to the drug file that includes binding affinity for specific receptors, leading to a much more robust opioid model.

### 3.2.4 Total Naloxone Administration

Despite the prevalence of naloxone accompanying emergency response providers, the rate of synthetic opioid overdoses and associated mortality is not decreasing. As stated above the use of multiple doses to reverse synthetic opiates is increasing in some localities, (Faul et al., 2017), although other studies show that in specific counties this trend is not mirroring the rise in opiate use, (Bell et al., 2019).

Generally, the use of naloxone has mirrored the rise in mortality in the age adjusted population, (Cash et al., 2018). This data points to a potent intervention tool that is potentially not being used correctly or not being sufficient to address the rise of synthetic opioids in the population.

To provide some understanding of the dosing requirements of naloxone to reverse extreme fentanyl exposure, we sought to provide a computational model that could start to answer that question.

Following the algorithm in Figure 4 we summed up the total naloxone reversal dose for a specific overdose event, Figure 10. For larger fentanyl doses, there seems to be an inflection point in the data, this occurs around the 0.9 mg dose and continues a rapid, nonlinear increasing response up to the highest range of fentanyl. We note that a standard dose of Narcan is 4mg, even though the standard dose we used for this analysis was 2mg, although some doses currently deployed are the 2mg formulation. We note that for a lower dose of synthetic opioids the standard dose of 2mg tends to reverse the respiratory depression sufficiently, with no relapse and with sufficient onset to deter the hypoxia seen the in the patient.

**Figure 4.**
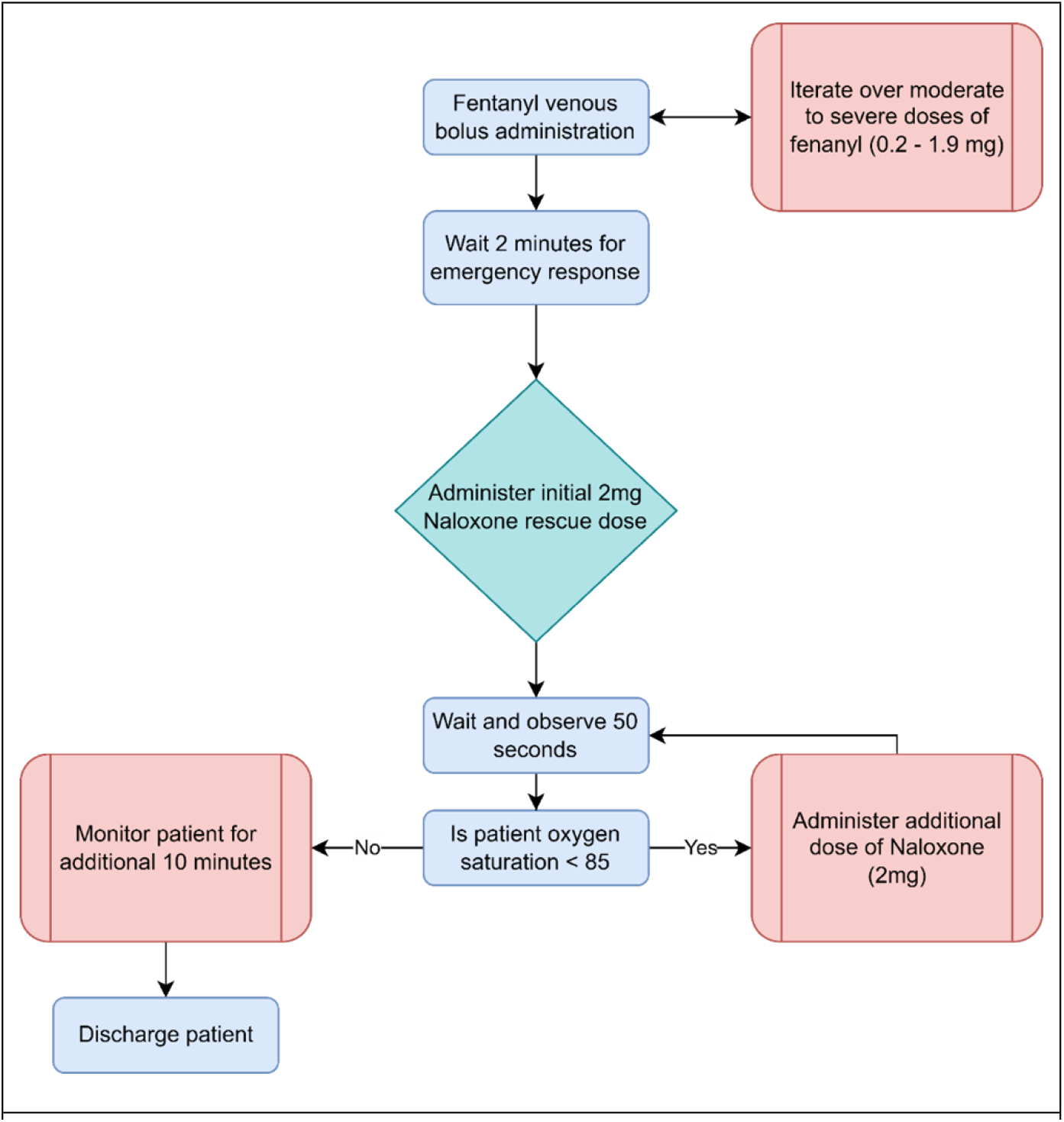
Simulated protocol for reversal resuscitation for an opioid overdose patient. Green denotes the “start” of the rescue period. Red boxes indicate decision/administration points, with blue boxes denoting observation or translation points.

**Figure 5.**
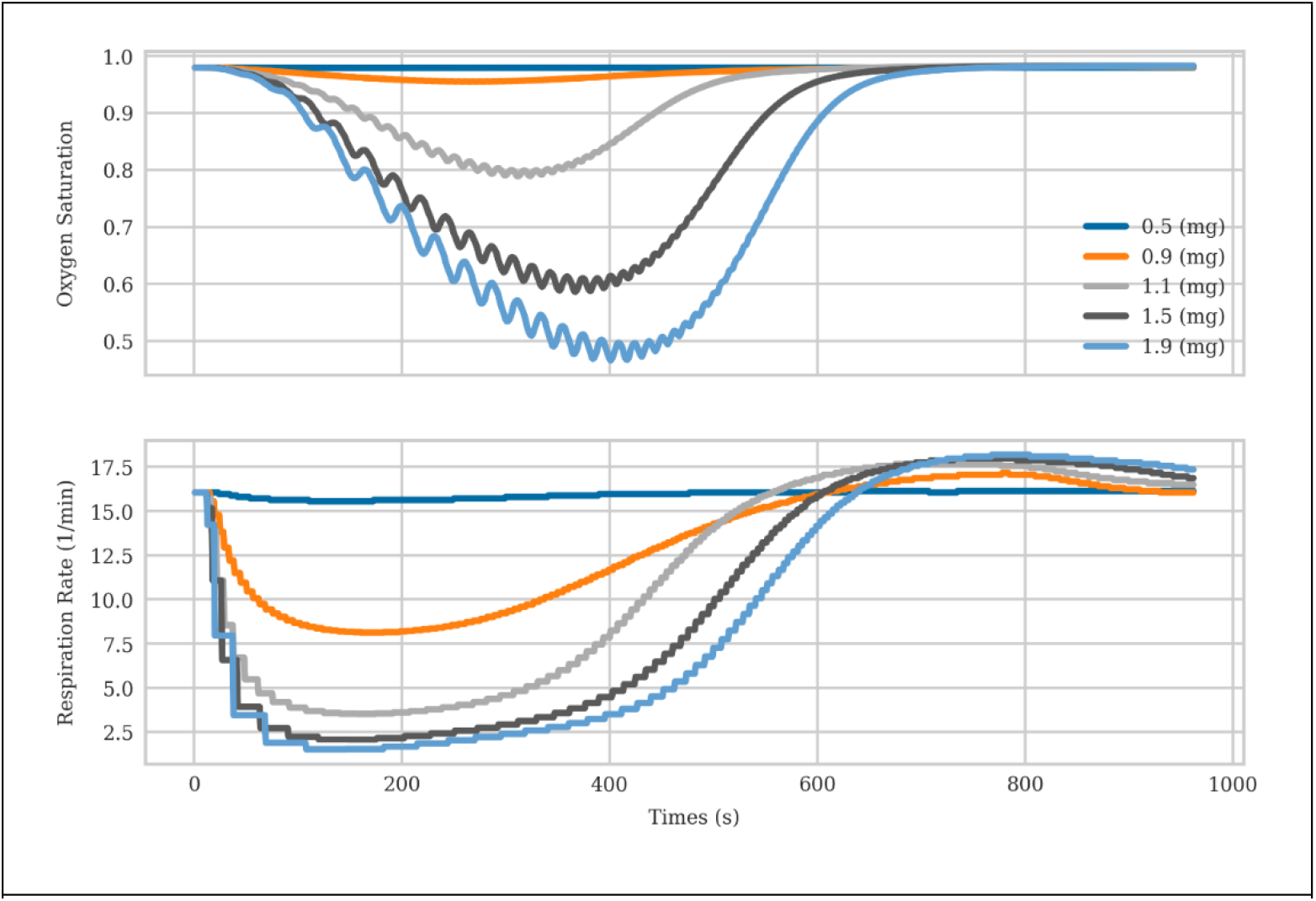
The primary cause and effect of fentanyl induced hypoxia are plotted here for varying levels of initial opioid dose. Oxygen saturation (top) is incurred through respiratory rate (bottom) depression.

**Figure 6.**
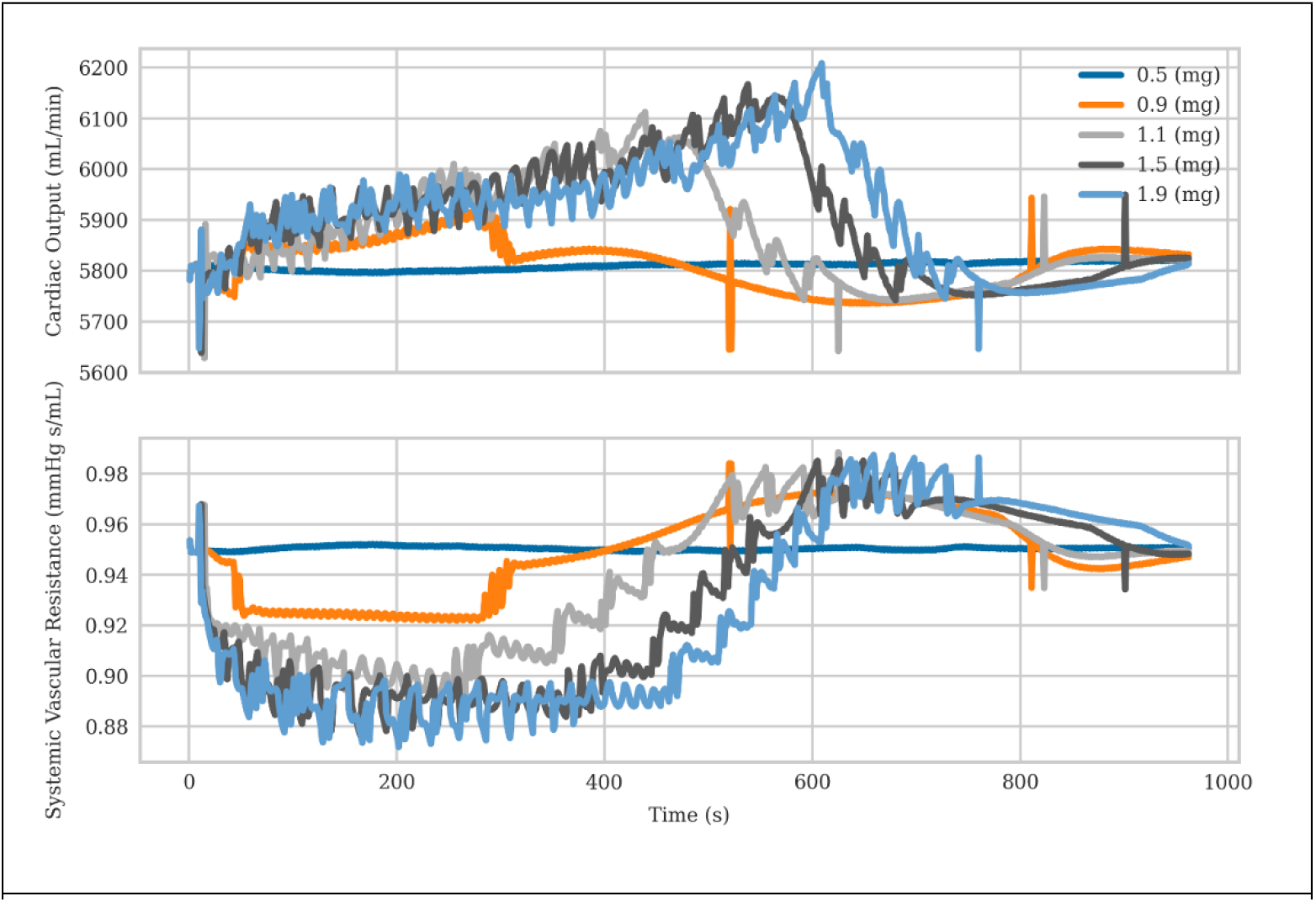
Cardiac output and systemic vascular resistance seen for varying levels of fentanyl administration. Rising cardiac output tries to compensate for the reduction in oxygen content in the blood.

**Figure 7.**
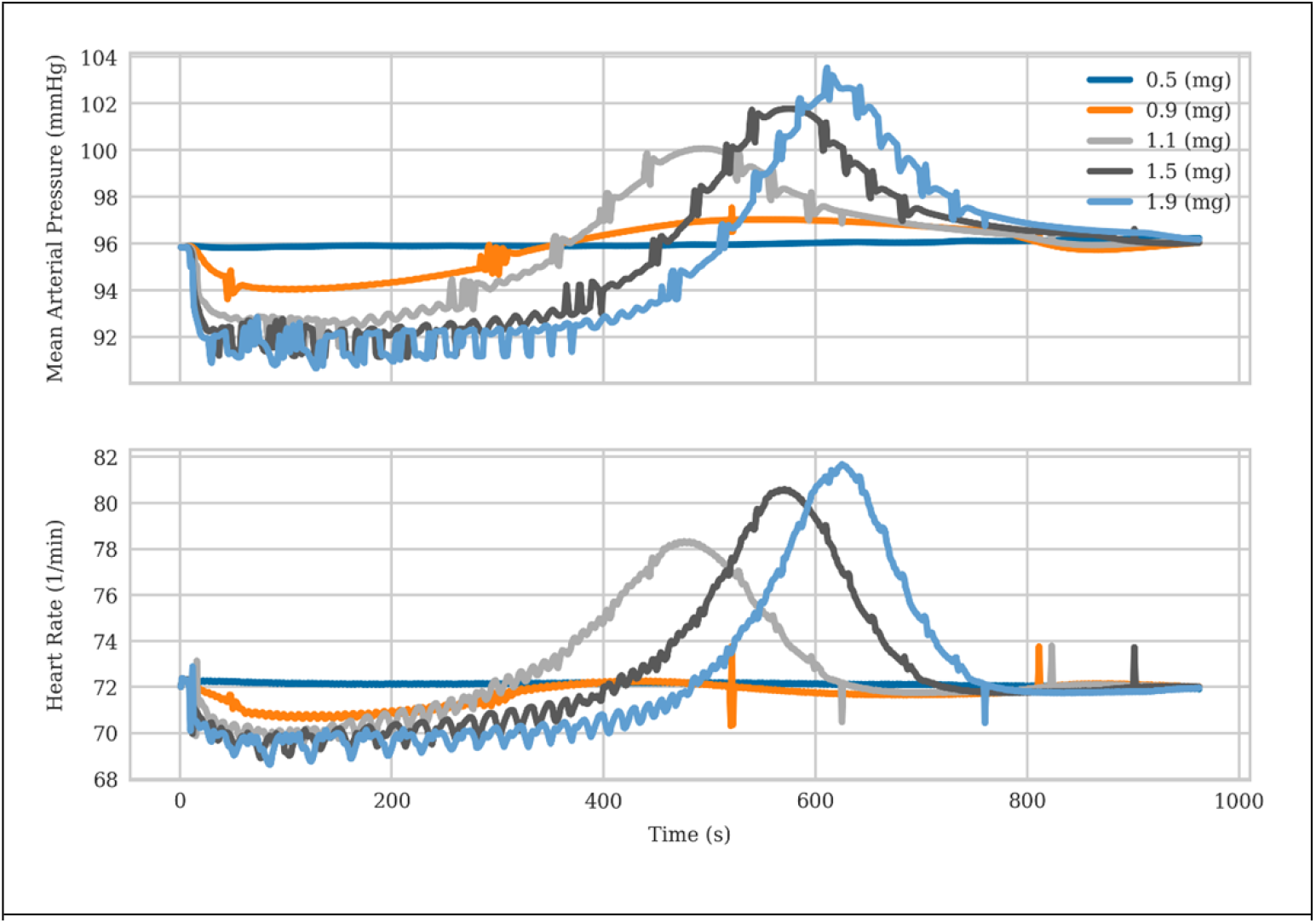
Mean atrial pressure (top) and heart rate for varying levels of opioid administration in the patient. Initial depression is due to the opioid with a recovery period following clearance and reversal.

**Figure 8.**
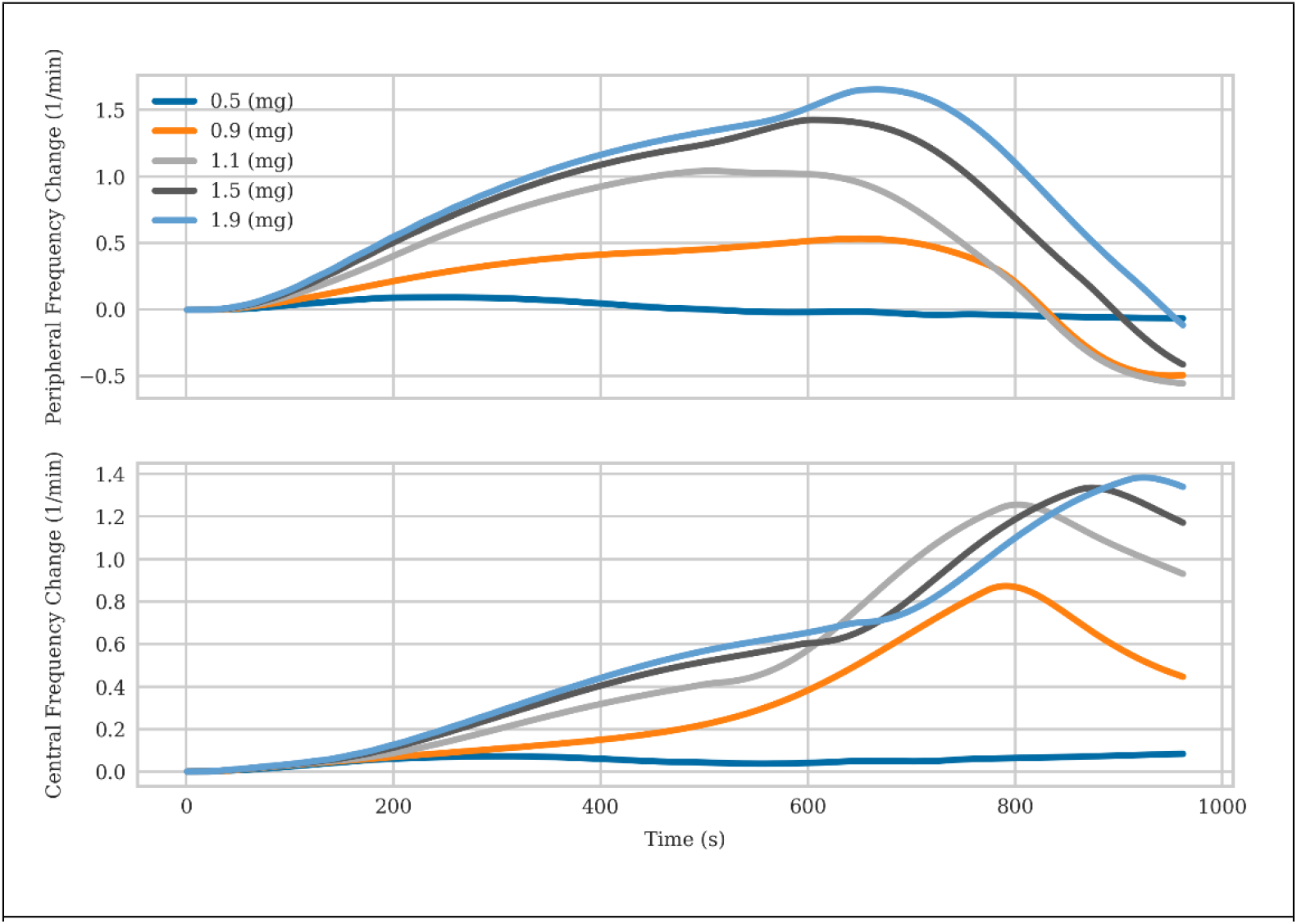
Central frequency function is reduced during the initial onset of the opioid. Because the opioid only targets the central nervous system, the peripheral frequency responds to the overdose event.

**Figure 9.**
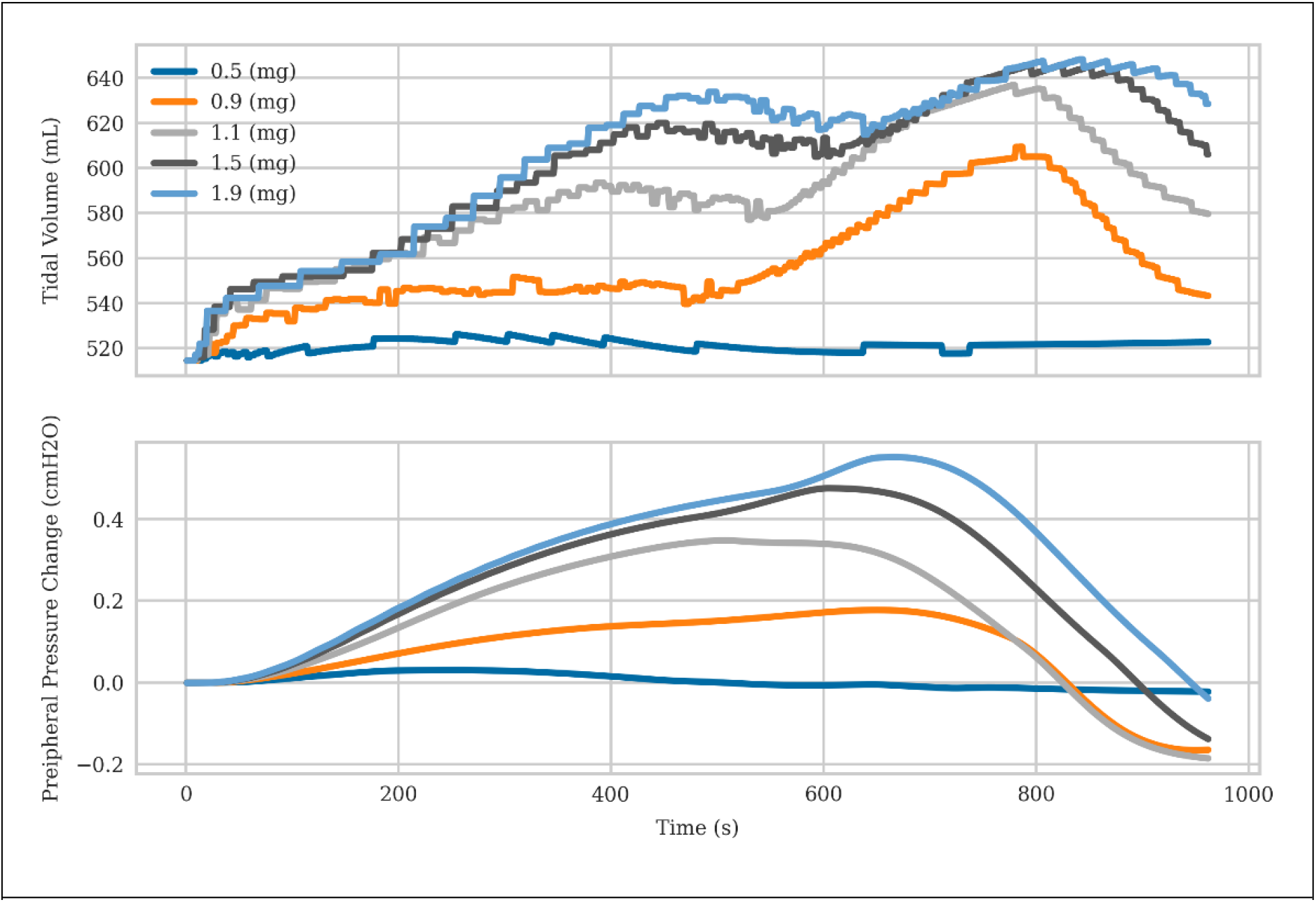
Peripheral pressure changes (bottom) react to the reduction in oxygen and correspondingly provide changes to the respiratory driver pressure, leading to an increase in tidal volume (top).

**Figure 10.**
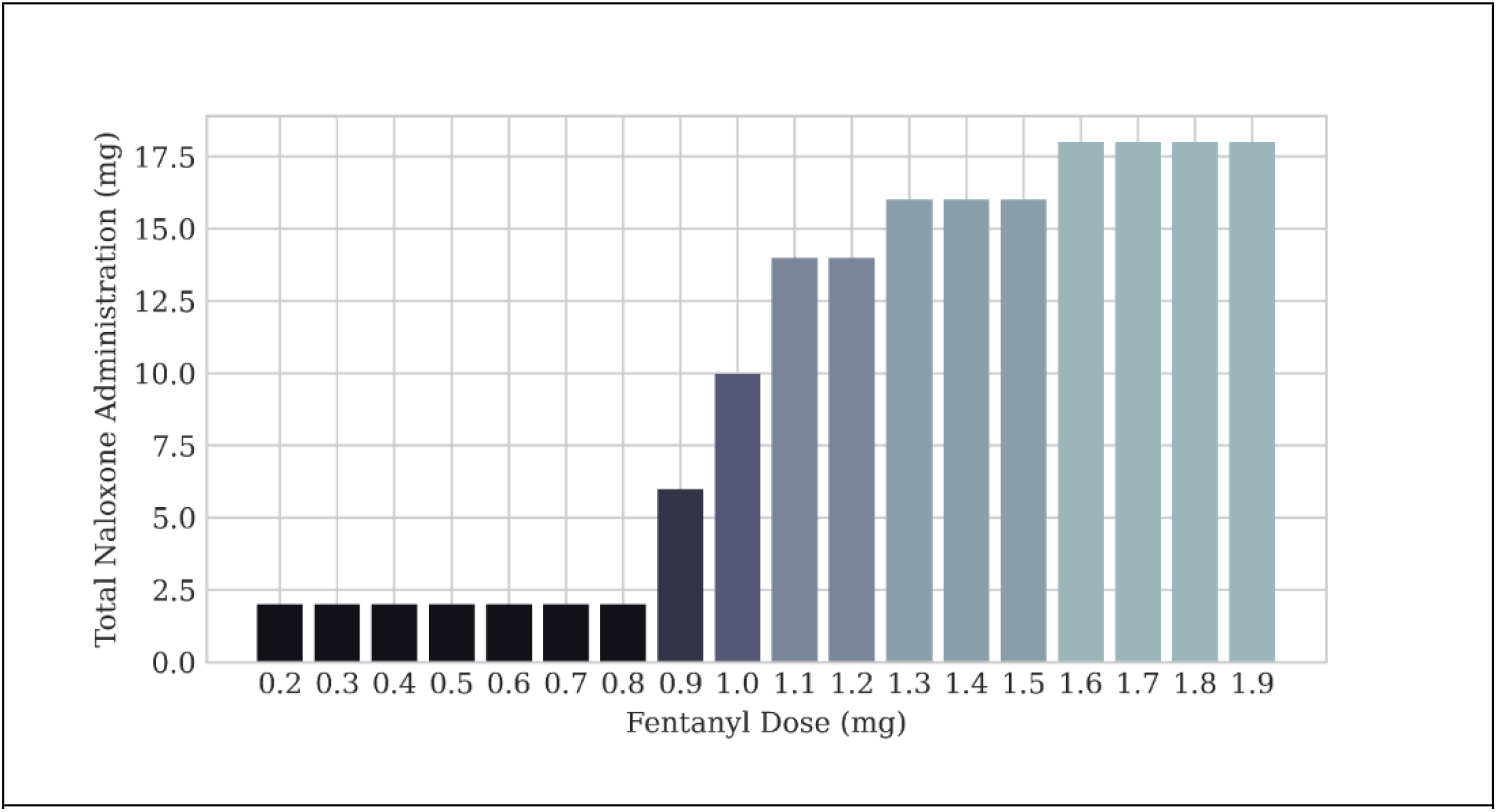
Total rescue dose administered for each fentanyl dose. A nonlinear response is seen as we transition from 0.8 to 1.1 mg amounts. Bars are colored as a function of total naloxone dose.

Noting that there existed an inflection point in the data, we sought to understand the general statistics between three distinct bins of fentanyl dose. We collected 6 doses and computed their mean, median and standard deviation, Figure 11. From these statistics we computed the p-value to compare whether each bin of fentanyl doses was statistically significant from the prior bin, Table 5. For each comparison we note the statistically significant dose of naloxone to reverse the opioid respiratory depression was required, confirming our hypothesis that the dose requirements for naloxone, via nasal administration, were insufficient for high levels of synthetic opioids.

**Figure 11.**
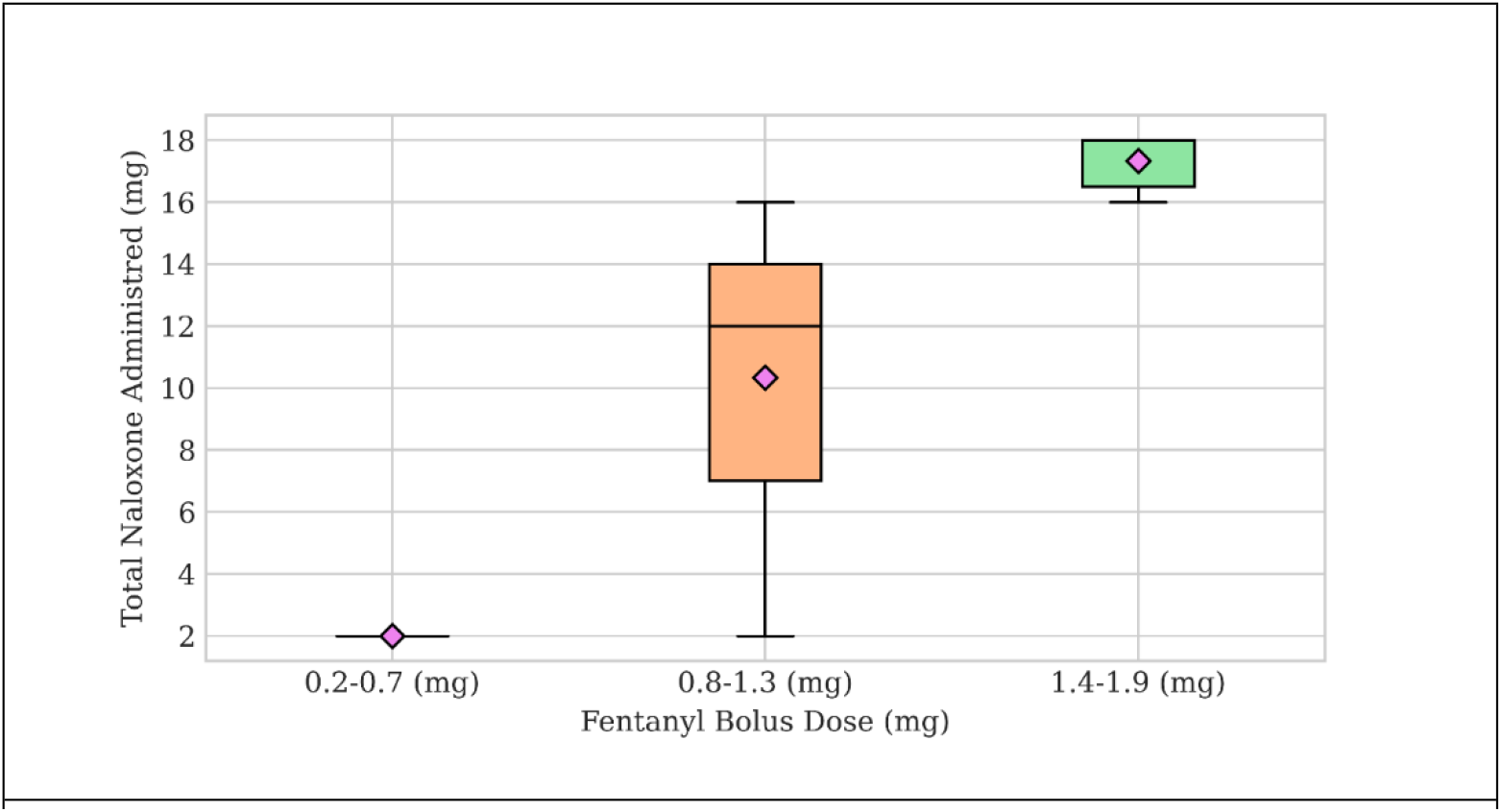
We group three fentanyl doses from analyzing the total administration amounts of the simulated experiment and compute the mean and standard deviation of the total naloxone dose. Each box plot quantifies the non-linear response of the required naloxone dose as a function of fentanyl dose.

**Table 5.** p-values reported for average naloxone dose required between the given fentanyl ranges reported. Values calculated via the medcalc online utility.

| Fentanyl Range Comparison | p-value |
| --- | --- |
| 0.2-0.7 v 0.8-1.3 | 0.0006 |
| 0.8-1.3 v 1.4-1.9 | 0.0018 |

## 4. Discussions

The opiate overdose death rate has surpassed motor vehicle collisions as the leading cause of death for ages 18-64 and for every death there are 6.4-8.4 non-fatal overdoses. The number of synthetic opiates related deaths is climbing and despite adequate supply of naloxone provided to first response providers, this death rate continues to climb (Skolnick, 2022; Zibbell et al., 2019). There are misconceptions and a general lack of knowledge surrounding the physiology of synthetic derived opiates which may be leading to under and ineffective utilization of naloxone at doses adequate to reverse the extremely high serum concentrations that these synthetic opiates reach at an impossibly short time interval, making them particularly lethal. These models quantify the dosing curves of the very high doses of fentanyl and the associated nasal naloxone treatment that are achieved in the blood which will be instrumental in understanding appropriate reversal dosing.

These models demonstrate previously held understandings that in higher doses of synthetic opiates, higher doses of naloxone should be administered to reverse the respiratory depression effects, up to 8x the standard dosing given by EMS. There is evidence that the timing of this administration, as seen in these models, also plays a role. One of the reasons synthetic fentanyl leads to such a frequency of overdose is the time to onset of respiratory depression. This type of rapid respiratory distress can be seen in the figures generated during this simulated treatment scenario. The peak concentration of opiate is higher and reached faster than seen previously in heroin overdose, 2 compared to 30 minutes (Vahedi et al., 2019). This degree of opiate intoxication leads to faster, more profound hypoxemia due to a decreased respiratory drive with a refractory recovery period compared to previously seen morphine equivalents. This leaves gaps for first responders regarding how many doses of naloxone may be required to adequately reverse an overdose.

Combining the physiologic effects of fentanyl with the timing of peak concentrations of the higher doses of naloxone can guide timing of monitoring after administration. Typically, if there is concern for severe opiate overdose, a patient should be transported to the nearest capable emergency department for monitoring and any further treatment that should be needed. In cases of refusal of transport, which occurs in approximately 40% of naloxone administration cases, patients requiring multiple doses of naloxone for reversal would imply higher serum concentrations of opiate and therefore a slower response for a sustained safer respiratory rate an oxygen saturation (Zozula et al., 2022). A higher dose of the opiates, as seen in these models, would also theoretically lead to a slower resolution of hypoxemia and bradypnea even with a normal concentration of naloxone dosing.

Understanding these physiologic derangements can offer guidance to first responders to not only the appropriate dosage but the time interval for recovery that should be anticipated.

The advantages of an accurate model of the pharmacokinetics of fentanyl, naloxone and associated respiratory depression are the ability to design and investigate repeat dosing scenarios in a safe environment. Limitations exist in animal models related to the translation of human to animal dosing equivalents, as well as obvious anatomical differences that may impact investigations regarding the route of Nalxone administration. Human models are limited to extremely low doses of opioids to ensure the safety of the participants of the study. A computational platform provides a safe, repeatable environment to design investigative studies as a first pass at obtaining information regarding changes in requirements of medical care. These investigations may also be a platform to inform future clinical trials and experimental studies that focus on naloxone plasma concentrations as a function of hypoxemia of the patient. How does naloxone interact with these opioids in the nervous system and what rate does it compete with these synthetic opioids as an agonist for selective nervous system binding. Including multiple receptor sites and associated impact on nervous function rather than just the mu pathway is one way to expand the current model.

Many flaws exist in the current model that need to be addressed in future iterations. These include nasal anatomical structure refinements, fentanyl overdose clearance and binding changes and nervous system model changes to consider the different binding pathways that the opioid may prefer depending on the type and amount administered. Clear issues regarding tidal volume compensation due to hypoxemia, seen at low doses of fentanyl, need to be addressed for higher levels of the opioid.

## 5. Conclusions

In conclusion, we present a model of the diffusion and transport of naloxone into the blood stream from a nasal administration pathway, coupled to a venous administration pathway of fentanyl. For all the drugs we compute their perfusion limited diffusion into the tissue spaces during transport, leading to clearance in the body. We demonstrate that the model of naloxone for high levels of administration validates well against experimental data. We couple the results of the plasma concentration of fentanyl to a pharmacodynamic model of the respiratory depression and mu receptor binding influence in a central nervous system model. We demonstrate that these coupled models can induce severe respiratory depression and associated hypoxemia because of high levels of fentanyl administration. In addition, we capture the reversal of the patient physiology due to the naloxone in the body due to a competitive antagonist drug-drug interaction model. We can see statistical significance due to the initial fentanyl dose and naloxone required to reverse the respiratory depression in the patient over time. We hope that this modeling result will induce more discussion regarding naloxone dosing requirements when it comes to synthetic opioid overdose for first responders in the field.

## 7. Author Contributions

The Author Contributions section is mandatory for all articles, including articles by sole authors. If an appropriate statement is not provided on submission, a standard one will be inserted during the production process. The Author Contributions statement must describe the contributions of individual authors referred to by their initials and, in doing so, all authors agree to be accountable for the content of the work. Please see here for full authorship criteria.

## 9. Acknowledgments

We’d like to acknowledge the support and guidance of Hugh Connacher, Harvey Magee Dr. Brett Talbot, and Geoff Miller who all provided excellent guidance during the original development of the BioGears project.

## 1. Data Availability Statement

The datasets generated and analyzed for this study can be found in the repository of the data and the python notebooks used to generate all images LINK. The software used to generate the provided data is also available free and open source in the BioGears core repository LINK, specifically the HowTo-NaloxoneNasal.cpp file was used to generate all the data in this paper.

